# Convergence of Ras- and Rac-regulated formin pathways is pivotal for phagosome formation and particle uptake in *Dictyostelium*

**DOI:** 10.1101/2022.06.21.497014

**Authors:** Sarah Körber, Alexander Junemann, Christof Litschko, Moritz Winterhoff, Jan Faix

## Abstract

Macroendocytosis comprising phagocytosis and macropinocytosis are actin-driven processes regulated by small GTPases that depend on the dynamic reorganization of the membrane that protrudes and internalizes extracellular material by cup-shaped structures. To effectively capture, enwrap, and internalize their targets, these cups are arranged into a peripheral ring or ruffle of protruding actin sheets emerging from an actin-rich, non-protrusive zone at its base. Despite extensive knowledge of the mechanism driving actin assembly of the branched network at the protrusive cup edge, which is initiated by the actin-related protein (Arp) 2/3 complex downstream of Rac signaling, our understanding of actin assembly in the base is still incomplete. In the *Dictyostelium* model system, the Ras-regulated formin ForG was previously shown to specifically contribute to actin assembly at the cup base. Loss of ForG is associated with a strongly impaired macroendocytosis and a 50% reduction of F-actin content at the base of phagocytic cups, in turn indicating the presence of additional factors that specifically contribute to actin formation at the base. Here, we show that ForG synergizes with the Rac-regulated formin ForB to form the bulk of linear filaments at the cup base. Consistently, combined loss of both formins virtually abolishes cup formation and leads to severe defects of macroendocytosis, emphasizing the relevance of converging Ras- and Rac-regulated formin pathways in assembly of linear filaments in the cup base, which apparently provide mechanical support to the entire structure. Remarkably, we finally show that active ForB, unlike ForG, additionally drives phagosome rocketing to aid particle internalization.

**Significance Statement:** Cup formation in macroendocytosis is a decisive, actin-dependent process that relies on distinct actin assembly factors generating the necessary mechanical forces to drive rearrangements of the plasma membrane and engulfment of extracellular material. Hitherto, in *Dictyostelium* the Arp2/3 complex and VASP were shown to promote actin assembly at the protrusive rim of phagocytic cups, while the Ras-regulated formin ForG generates about half of the actin filament mass at the base. Here, we show that ForG synergizes with the Rac-regulated formin ForB to form the bulk of filaments at the cup base. Loss of both formins virtually abolishes cup formation and leads to dramatic defects in macroendocytosis, illustrating the relevance of converging Ras- and Rac-regulated signaling pathways in this process.

## Introduction

Macroendocytosis comprises phagocytosis and the closely related macropinocytosis, in which larger particles or bulk fluid are taken up from the extracellular space by actin-rich, cup-shaped cell protrusions, respectively (1, 2). In higher eukaryotes, phagocytosis by so-called professional phagocytes of the innate immune system (3) is crucial for immune surveillance and tissue homeostasis by clearance of pathogens, apoptotic cells or debris (4, 5), whereas in lower eukaryotes such as the soil-dwelling amoeba *Dictyostelium discoideum*, phagocytosis is primarily used for nutrient acquisition (6, 7). Therefore, it has been widely assumed that the evolutionary origin of phagocytosis is the uptake of nutrients by simple unicellular eukaryotes (8, 9). Moreover, phagocytosis has been in the focus of lively discussions of eukaryogenesis for many years, where it has been argued that it is either a requirement for or a consequence of the acquisition of the ancestral mitochondrion (9). Despite serving different purposes in higher and lower eukaryotes, the regulation and the core mechanisms of phagocytosis and macropinocytosis driven by actin polymerization are highly conserved across species (9, 10). Given its repertoire of actin-binding proteins and regulators of the actin cytoskeleton that is comparable to that of mammalian cells (11, 12) as well as its ease of manipulation and availability of laboratory strains with highly upregulated rates of macropinocytosis (13, 14), *Dictyostelium* has proven an excellent system for the study of macroendocytosis serving as a model for immune cells (15).

Although macropinocytosis and phagocytosis are mechanistically similar processes that share many components with the machinery driving protrusion of lamellipodia or pseudopodia in motile cells (16), there are nevertheless important differences between the two internalization pathways (2). Phagosomes are typically sculpted by a receptor-guided, zipper-like advance of the membrane and the cytoskeleton over the particle they ingest (2), whereas the structurally related macropinosomes can form autonomously in the absence of any external guidance cues (8, 17). Macropinocytic cups appear to be organized around a signaling patch of the phosphoinositide PIP3, active Ras and active Rac that guides actin polymerization to its periphery, whereas phagocytic cups are initiated primarily by receptor activation and localized signaling due to contact with the prey (8). While in both cases active Rac is important for regulating actin protrusions, active Ras is restricted to the interior of the cup (17). The first important step of phagocytosis entails particle recognition and binding through a diverse set of receptors to initiate phagocytosis (18). The most extensively studied phagocytic receptors of mammalian cells include the Fcγ- and the integrin-type CR3 receptors, that interact with antibody-or complement-opsonized particles, respectively (18, 19). *D. discoideum* cells use the integrin-like receptors SibA and SibC for adhesion and phagocytosis whereas the G protein-coupled receptor fAR1 is used for both chemotaxis and phagocytosis (15, 20, 21). Receptor-binding prompts local actin assembly to initiate formation of a phagocytic cup, followed by extension of pseudopods wrapping around the particle, and finally terminates by the closure of the cup through fusion of converging protrusions followed by disassembly of the F-actin coat after particle internalization (2, 22).

Comparable to pseudopods and lamellipodia of motile cells, actin assembly at the tips of macroendocytic cups is initiated by the Arp2/3 complex downstream of SCAR (suppressor of cAMP receptor)/WAVE complex (also known as the WAVE-regulatory complex WRC) activation and Rho-family GTPase signaling (23). Consistently, inactivation of SCAR in *Dictyostelium*, which accumulates in a ring-like fashion along the cup edge (17), diminishes F-actin at the protrusive cup rim and is accompanied with significantly reduced rates of macropinocytosis and phagocytosis (24). The barbed ends of newly formed filament branches are eventually capped by heterodimeric capping protein (25). However, actin filament elongation factors such as Ena/VASP or formin family proteins protect filament ends from capping and also markedly accelerate filament elongation rates (26, 27). Similarly, elimination of the single Ena/VASP member in *Dictyostelium* cells VASP, which also accumulates as a ring at the protrusive cup rim, results in markedly reduced rates in macroendocytosis (28).

Formins are dimeric multidomain proteins that nucleate and elongate linear actin filaments (29) such as those found in filopodial bundles, the cleavage furrow or the cell cortex (30, 31). The proline-rich formin homology domain 1 (FH1) recruits profilin (PFN)-actin complexes to enhance filament elongation by the adjacent FH2 domain (32). Members of the subfamily of *Diaphanous*-related formins (DRFs) are tightly regulated. By virtue of regulatory sequences located in the N- and C-terminal regions, these proteins are intrinsically autoinhibited. Typically, the binding of Rho-subfamily GTPases to the N-terminal GTPase-binding domain (GBD) releases autoinhibition mediated by interaction of the C-terminal diaphanous-autoinhibitory domain (DAD) with the GBD, thus rendering the protein active and allowing it to interact with downstream effectors (32, 33). Like all Ras-family members, Rho-subfamily GTPases cycle between an active GTP-bound state and an inactive GDP-bound state to regulate various aspects of intracellular actin dynamics including macroendocytosis (34–37). In contrast to the Arp2/3 activator SCAR/WAVE, which was previously found to localize exclusively at the rim of macropinocytic cups, F-actin is highly enriched in the entire structure beneath the plasma membrane (38). Remarkably, in *D. discoideum* the Ras-regulated formin ForG was shown to specifically contribute to actin assembly at the non-protrusive base of macroendocytic cups (39). Loss of ForG is accompanied by a 50% reduction of F-actin content in the base of phagocytic cups and is associated with strongly impaired macroendocytosis (39), which additionally suggests the presence of other factors that specifically contribute to actin assembly in the cup base.

Here, we identify the Rac-regulated DRF ForB as an additional actin assembly factor prominently localizing to endocytic cups in its active form. Consistent with its rather weak actin assembly activity *in vitro*, individual loss of ForB does not noticeably interfere with macroendocytosis. However, the combined elimination of ForB and ForG virtually abolishes phagosome formation and leads to severe defects in macroendocytosis, illustrating the relevance of converging Ras- and Rac-regulated formin pathways in assembly of the cup base, which apparently serves as a mechanical support of the entire structure.

## Results

### ForB localizes to macropinosomes and phagosomes

Given that the actin signal at the base of cups in *forG*^*-*^ mutants is only diminished by about 50% and the cells still capable to internalize particles, albeit with a reduced rate (39), we reasoned that an as yet uncharacterized formin could synergize with ForG in cup formation. The *D. discoideum* genome encodes ten formins, from which five (ForA, ForB, ForE, ForG, ForH) are predominantly expressed during the growth phase where macroendocytosis occurs (40). Four of them have already been attributed specific cellular functions: ForG in macroendocytosis and ForA, ForE, and ForH in regulation of actin cell cortex mechanics (31), thus leaving ForB as the only remaining candidate. Although a previous study found no detectable null-mutant phenotype (41), more recent evidence suggests ForB to be involved in the propagation of actin waves (42), which resemble frustrated phagocytic cups, pointing towards a potential role of ForB in macroendocytosis.

Formin B (ForB) is a DRF with the characteristic domain architecture GBD/FH3-FH1-FH2-DAD (40). As expected, due to autoinhibition, ectopically expressed full-length ForB N-terminally tagged with GFP localized uniformly in the cytoplasm of Ax2 wild-type (WT) cells (Fig. 1*A*). Thus, we monitored localization of a constitutive active variant missing the C-terminal DAD domain (YFP-ForB-ΔDAD) and of the N-terminal region encompassing the GBD/FH3 domain alone (YFP-ForB-N) fused to YFP. Both constructs localized prominently to macropinosomes (Fig. 1*B* and *C*), although the N-terminal construct exhibited a higher nonspecific background (Fig. 1*B*). Thus, if not stated otherwise, in all subsequent localization studies, constitutively active YFP-ForB-ΔDAD construct was used. Co-expression of YFP-ForB-ΔDAD with the F-actin marker LifeAct fused to mRFP in growth-phase cells revealed ForB to co-localize with filamentous actin in ruffles and macropinosomes (Fig. 1*D*, *SI Appendix*, Movie S1). Confocal time-lapse imaging of YFP-ForB-ΔDAD in presence of TRITC-labelled yeast particles further revealed that ForB is also enriched along the entire actin-rich base of phagocytic cups throughout particle engulfment, but vanishes rapidly after cup closure (Fig. 1*E*, *SI Appendix*, Movie S2). Combined, these results corroborate further evidence for a functional role of ForB in macroendocytosis.

**Fig. 1.**
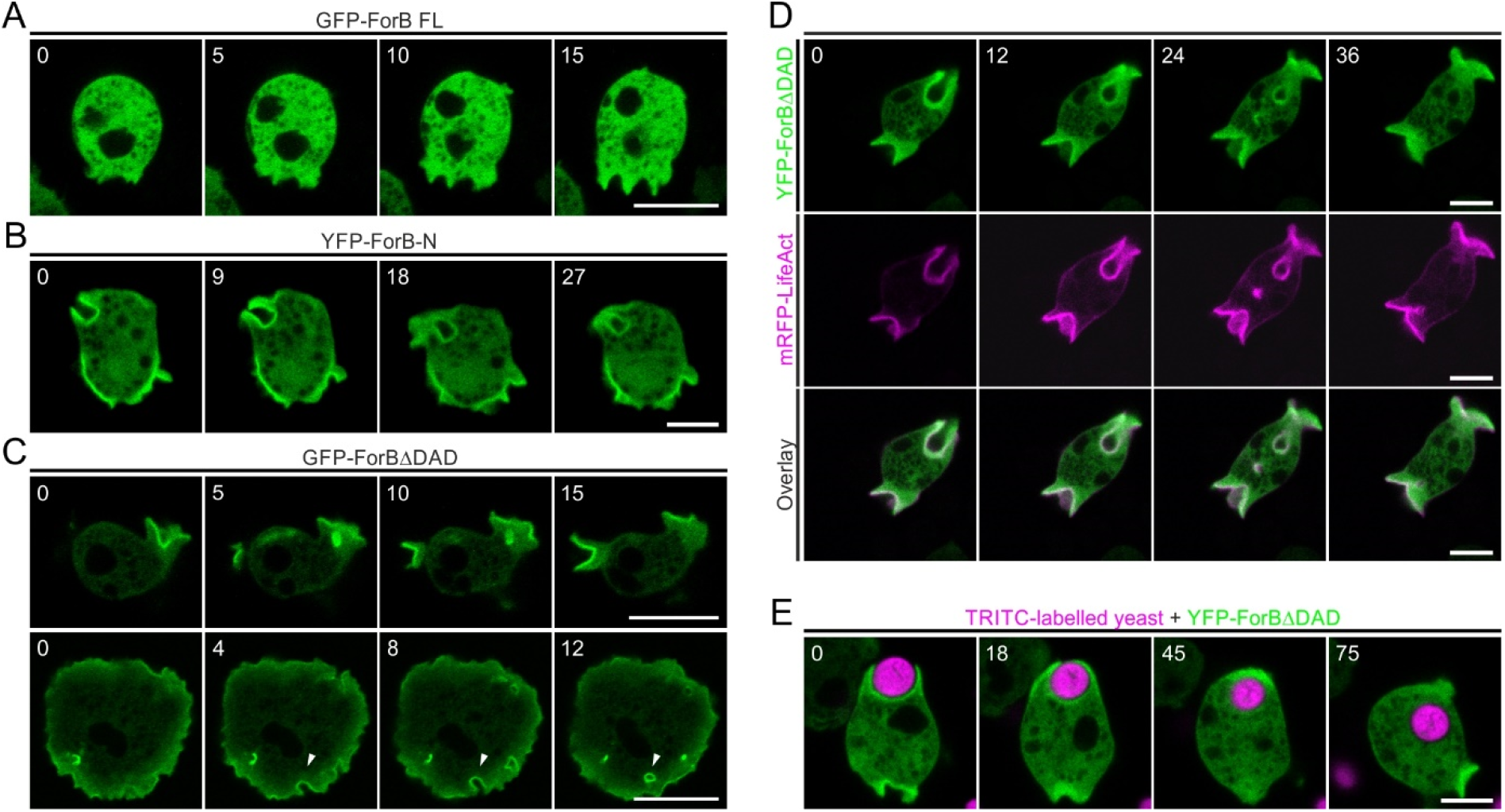
ForB accumulates in endocytic cups. (*A*) Representative images of autoinhibited, full-length (FL) ForB fused to GFP shows uniform cytosolic localization. (*B* and *C*) YFP-tagged ForB-N (residues 1-406) and constitutively active ForBΔDAD (residues 1-1,081) fused to GFP prominently localized to nascent macropinosomes and ruffles that converted into macropinosomes (white arrow heads). (*D*) Time-lapse imaging of WT cells coexpressing LifeAct-mRFP and YFP-ForBΔDAD revealed striking colocalization of ForB and F-actin at macropinosomes. (*E*) YFP-ForBΔDAD also accumulated at phagocytic cups throughout engulfment of large TRITC-labelled yeast particles. Confocal section are shown. Time in seconds. Scale bars, 5 μm (B, D and E) and 10 μm (A and C).

### Combined loss of ForB and ForG severely impairs phagocytosis

Next, we sought to validate expression of ForB during the vegetative stage by immunoblotting, since previous studies analyzed expression exclusively on RNA level. A developmental time course confirmed ForB to be expressed at similar levels during the vegetative state and the first 9 hours of development as assessed by the appearance of the early developmental marker contact site A (csA) (43) (Fig. 2*A*). To assess the function of ForB in macroendocytosis, we eliminated ForB in Ax2 WT cells by homologous recombination (Fig. 2*B*). To allow examination of a potential synergy of ForB with ForG in large-scale endocytosis we used the *forG*^*-*^ single mutant (39) and additionally disrupted the *forB* gene to obtain a double mutant devoid of both formins. Loss of ForB in single and double mutants was confirmed by immunoblotting (Fig. 2*B*). During isolation of the mutants, we noticed that the double mutant formed much smaller colonies on bacterial lawns as compared to control. Thus, we measured the rate at which cells form bacteria-free plaques when grown on a dense lawn of *Klebsiella aerogenes* as the only nutrient source. *Dictyostelium* cells cannot propagate on such agar plates, so their ability to grow in the presence of the bacteria, reflected in the size of the plaques that formed after seven days, is mainly due to their ability to phagocytose the bacteria. The *forB*^*-*^ single mutant formed plaques that were only slightly smaller as compared to the size of plaques produced by WT and *forG*^*-*^ mutant cells. However, the combined elimination of both formins resulted in plaques that were more than 94% smaller in surface area than those formed by WT cells (Fig. 2*C* and *D*), strongly suggesting a synergistic role of ForB and ForG in phagocytosis. We then examined the growth of WT and mutant cells in liquid growth medium in shaken suspension as a proxy for macropinocytosis. Unexpectedly, the growth curves of *forB*^*-*^ cells and WT cells were highly similar, with calculated mean generations times of 11.0±0.7 h and 11.1±1.0 h (mean±SD), respectively (Fig. 2*E*). The *forG*^*-*^ cells grew slightly slower compared to the WT and had a mean generation time of 11.7±0.7 h whereas the *forB*^*-*^/*forG*^*-*^ double mutant had with 12.8±0.5 h the slowest generation time. Then we directly monitored macropinocytosis by measuring the uptake of the fluid-phase marker TRITC-Dextran (70 kDa). Compared to the WT control, the relative uptake rate in *forB*^*-*^ cells dropped by 13%, while it dropped by 29% in *forG*^*-*^ cells (Fig. 2*E*). This notwithstanding, the *forB*^*-*^/*forG*^*-*^ double mutant showed even a slightly lower uptake rate compared to *forG*^*-*^ cells.

**Fig. 2.**
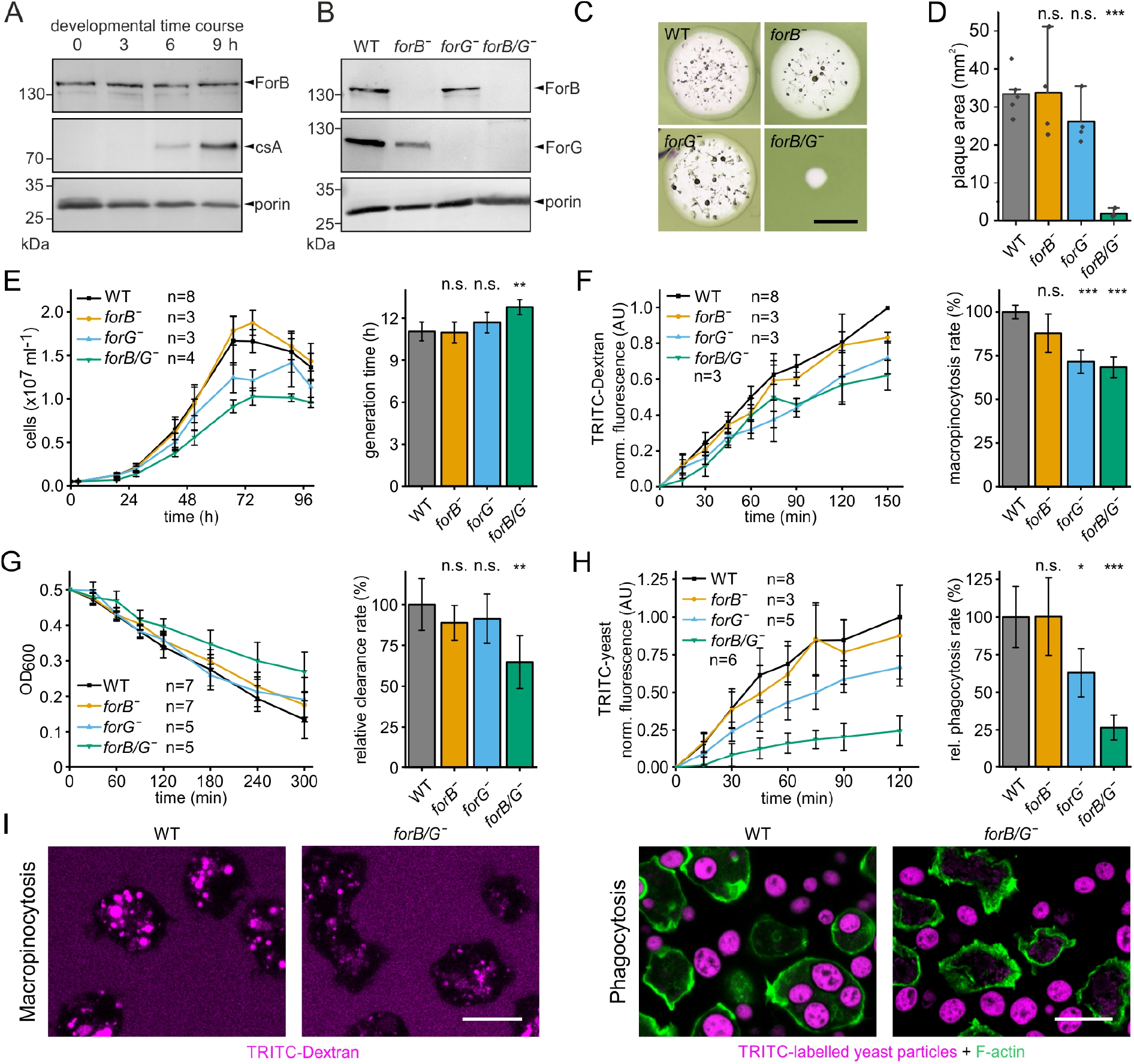
Combined loss of ForB and ForG dramatically impairs phagocytosis. (*A*) Expression profile of ForB during early development of *D. discoideum* WT cells. Time of starvation is indicated in hours (h). Contact site A (csA) served as early aggregation marker and porin was used as loading control. (*B*) Loss of ForB and ForG in single and double mutants was confirmed by immunoblotting using specific antibodies. Porin was used as loading control. (*C*) Representative plaques formed by WT and mutant cells indicated on bacterial lawns of *Klebsiella aerogenes* 168 h post seeding. Scale bar, 3 mm. (*D*) Quantification of plaque surface area after 168 h. Mean±SD. Pooled data from 4-5 independent experiments with >10 plaques each are shown. ** p≤0.01, n.s. not significant. (*E*) Growth curves of WT and mutants cells in shaken suspension and derived generation times. (*F*) Quantification of macropinocytosis by the uptake of TRITC-Dextran (70 kDa) and derived rates. (*G*) Clearance of bacteria from shaken suspension revealed the strongest defect in the *forB*^*-*^/*forG*^*-*^ double mutant. (*H*) Quantification of phagocytosis using TRITC-labelled yeast particles. (*I*) Accumulation of TRITC-dextran (magenta) by macropinocytosis after 60 min of incubation (left) and of TRITC-labelled yeast particles (magenta) by phagocytosis after 40 min of incubation (right) visualized by confocal microscopy. WT and double-mutant cells are visualized by F-actin staining (green). Scale bars, 10 μm. Mean±SD displayed for all experiments, n: number of independent experiments. * p≤0.05, ** p≤0.01, *** p≤0.001, n.s. not significant.

Next, we quantified clearance of bacteria from shaken suspension by phagocytosis. Consistent with the formation of smaller plaques on bacterial lawns, we observed a markedly diminished clearance rate of bacteria only upon combined loss of ForB and ForG (64.8%±16.2 of WT), whereas the single knockout mutants displayed slightly lower, albeit not statistically significant rates of about 90% as compared to the WT (Fig. 2*G*). Notably, the defect in phagocytosis was even more pronounced during uptake of the much larger yeast particles, with the *forB*^*-*^/*forG*^*-*^ mutant having an uptake rate of only 26.5±8.3% compared to the WT (Fig. 2*H*). Consistently, once again, elimination of ForB alone did not lead to a defect (100.3±25.8%) in uptake, whereas loss of ForG reduced the uptake rate to 62.9±16.0% compared to the WT, which is in excellent agreement with previous work (39). Taken together, these data clearly show that ForB contributes to macropinocytosis, but primarily to phagocytosis, although the consequences of its loss become evident only in the concurrent absence of ForG (Fig. 2*I*).

### ForB is a weaker actin assembly factor compared to ForG

Thus, to better understand the different contributions of these formins to macroendocytosis, we compared the biochemical activities of ForB and ForG *in vitro*. Since expression of a C-terminal ForB fragment harboring the complete FH1 domain with its four highly repetitive poly-proline stretches was not possible in *E. coli*, we purified a ForB fragment with two poly-proline stretches (ForB-2P) preceding the FH2 and DAD domains for biochemical analysis. Bulk polymerization assays with pyrene-actin and increasing amounts of ForB-2P in the higher nanomolar range revealed that ForB stimulated spontaneous actin assembly in a concentration-dependent manner (*SI Appendix*, Fig. S1*A*), albeit with substantially lower activity compared to a corresponding C-terminal ForG fragment (ForG-3P), which has been characterized previously (39) and was used as a reference. Subsequently, both formin fragments were analyzed for nucleation activity by total internal reflection fluorescence microscopy (TIRF-M) at the single-filament level. A relatively high concentration of 250 nM ForB-2P had to be used to obtain a visible and quantifiable increase in nucleation, and once again, it was almost 3 times less effective compared to ForG-3P (*SI Appendix*, Fig. S1*B* and *C*). Quantification of filament elongation in the presence of *Dictyostelium* profilin I (PFN I) revealed that ForB-2P with 16.9±2.2 subunits/s only slightly exceeded the growth of control actin control filaments with 12.8±2.1 subunits/s, while ForG-3P reached elongation rates of 51.7±3.2 subunits/s (*SI Appendix*, Fig. S1*C*). These data not only show that, at least *in vitro*, ForB is a relatively weak actin assembly factor compared to ForG, but may also explain the lack of a detectable phenotype in the single knockout mutants by the presence of the more powerful formin ForG.

### ForB interacts with active RacB

ForB encompasses a canonical GTPase-binding domain (GBD) characteristic for DRFs (40). We therefore explored which GTPase could be responsible for activation of ForB by yeast two-hybrid (Y2H) analyses, using a N-terminal ForB fragment (ForB-N) harboring the GBD and the FH3 region (amino acids 1-406) as bait. *Dictyostelium* cells lack genuine Rho and Cdc42 homologs, but express 20 Rac-subfamily GTPases (44). Thus, we systematically screened ForB-N with all these Rac GTPases in their constitutively-activated forms derived by amino-acid substitutions corresponding either to position 12 within the P-loop or to position 61 within switch II of human Rac1, respectively (37). Given that ForG was previously found to interact with the Ras-family proteins RasB, RasG and RasD (39), we additionally tested active variants of all 13 Ras-family GTPases as bait in the Y2H screen. ForB did not interact with any of these Ras GTPases, except with RasB Q61L, which, however, proved to be nonspecific as growth of yeast cells on selective media was even observed in the absence of the bait (*SI Appendix*, Fig. S2*A*). In contrast, ForB interacted with several active members of the Rac family under medium-strength screening conditions including Rac1a-c, RacB, RacF1, RacH, and RacM, albeit interactions with RacH and RacM also turned out to be false-positive (*SI Appendix*, Fig. S2*B*) (39). Notably, upon selection at highest stringency, ForB interacted only with the active RacB variants G12V and Q61L, but not with the dominant-negative form T17N, thus rendering this GTPase the most likely activator of ForB (Fig. 3*A*).To validate the ForB-RacB interaction in an independent assay, we employed bimolecular fluorescence complementation (BiFC) that enables direct visualization of protein-protein interactions in living cells (45). To this end, ForB-ΔDAD was fused to the C-terminal fragment of mVenus and interactions were tested in combination with either dominant-negative RacB T17N, RacB WT, or the constitutively active variant RacB Q61L fused to the N-terminal fragment of mVenus. Of note, fluorescence complementation was only seen with WT as well as constitutively active RacB, but not with dominant-negative RacB that served as negative control (Fig. 3*B*). Moreover, the fluorescence signal formed with RacB WT was distributed along the entire periphery of the cell, whereas localization with the RacB Q61L variant was more confined to phagosomes, macropinosomes, ruffles, and protrusions. The excellent fluorescence and localization with the constitutively-active RacB variant allowed us to perform confocal, time-lapse imaging showing that the ForB-RacB complex is specifically restricted to phagosomes and macropinosomes (Fig. 3*C* and *D*, *SI Appendix*, Movies S3 and S4). Combined, these data demonstrate that active RacB also specifically interacts with ForB *in vivo*.

**Figure 3.**
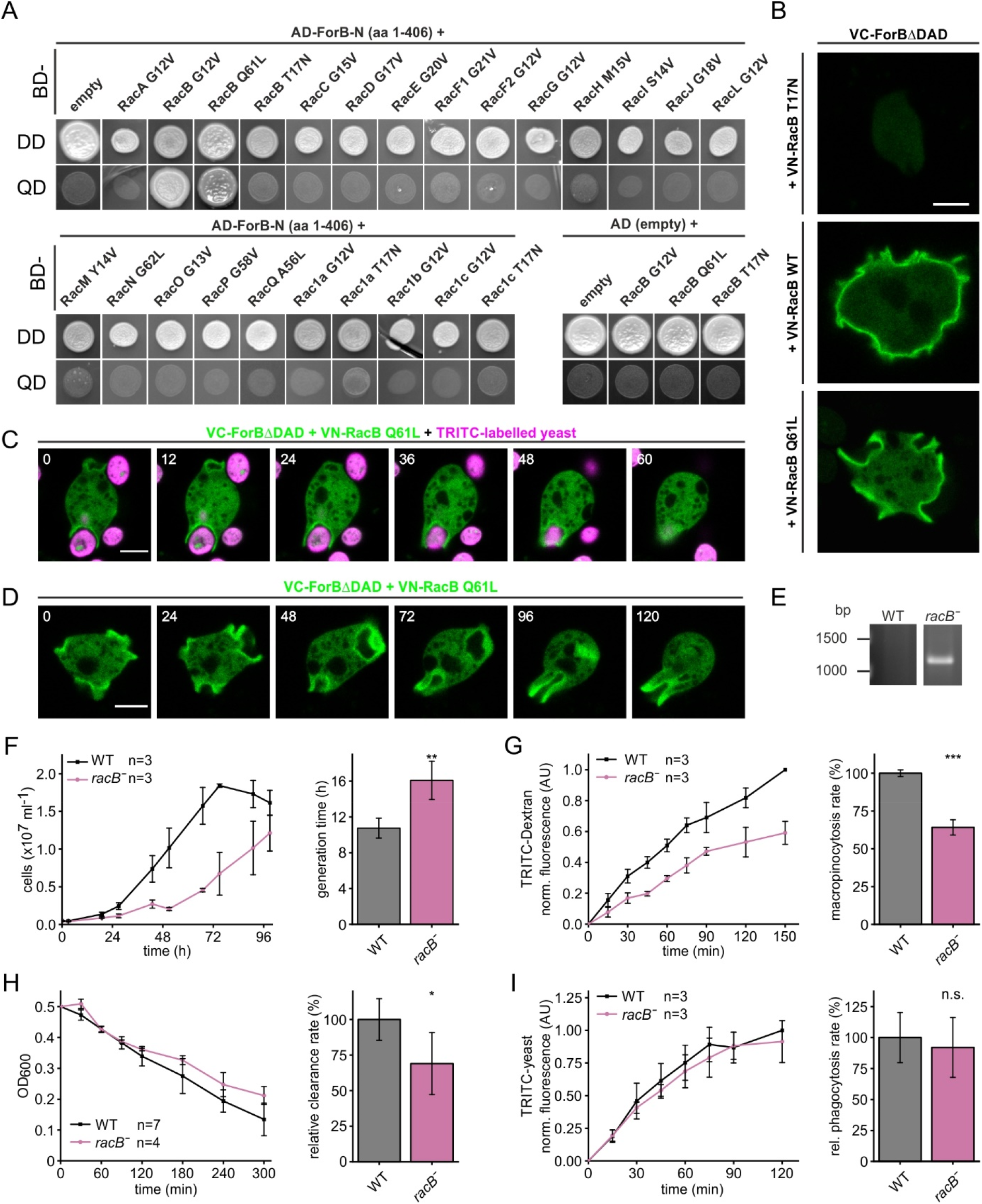
ForB interacts with active RacB. (*A*) Y2H screen of ForB-N with all 20 Rac GTPases of *D. discoideum* showed strongest interaction with constitutively-active RacB. The presence of bait and prey plasmids was scored by growth by double-dropout media (DD) lacking leucine and tryptophan and genetic interactions were tested by growth at highly stringency on quadruple-dropout media (QD) lacking leucine, tryptophan, adenine and histidine. Positive hits were additionally validated in screens using respective dominant-negative variants and empty AD plasmids as negative controls. AD, Gal4-activation domain; BD, Gal4-binding domain. (*B*) Bifluorescence complementation (BiFC) confirmed interaction of ForB-ΔDAD with constitutively-active RacB *in vivo*, whereas no interaction was found with dominant-negative RacB (T17N). Scale bar, 5 μm. VC, mVenus C-terminal fragment; VN, mVenus N-terminal fragment. Still images of confocal, time-lapse movies illustrating WT cells expressing BiFC constructs with constitutively active RacB Q61L during phagocytosis (*C*) or macropinocytosis (*D*). Scale bars, 5 μm. (*E*) Disruption of the *racB* gene was validated by screening for the insertion of the Blasticidin-resistance cassette using PCR. (*F*) Growth of *racB*^*-*^ cells in shaken suspension was severely impaired and markedly increased generation time. Impaired uptake of TRITC-dextran by macropinocytosis (*G*) and clearance of bacteria (*H*) of *racB*^*-*^ cells. (*I*) Quantification of phagocytosis with TRITC-labeled yeast revealed uptake of particles was not substantially affected. F-I, mean±SD. n, number of independent experiments. * p≤0.05, ** p≤0.01, *** p≤0.001, n.s. not significant.

To explore whether loss of RacB would mimic the defects of *forB*^*-*^ in macroendocytosis, we generated a *racB*^*-*^ mutant by homologous recombination (Fig. 3*E*), and assayed the mutant for a number of parameters in large-scale endocytosis. In comparison to the WT, the *racB*^*-*^ mutant grew considerably slower in shaken suspension containing growth medium and exhibited markedly diminished pinocytosis rates (Fig. 3*F* and *G*). The uptake of bacteria and large yeast particles was reduced by 30% and 8%, respectively (Fig. 6*H* and *I*). Thus, although inactivation of the main regulator RacB partially phenocopies the defect of the *forB*^*-*^ cells, it has a stronger effect on large-scale endocytosis. This notwithstanding, RacB is expected to regulate and activate additional downstream targets such as the WRC and Arp2/3 complex, which also drive macroendocytosis.

### ForB^-^/forG^-^ mutants exhibit a dramatically reduced F-actin content in the phagosome base

To assess potential defects in actin assembly during large-scale endocytosis, we then monitored and compared the F-actin distribution in semi-closed phagosomes of WT and mutant cells engulfing fluorescent yeast particles after phalloidin staining. Line scan measurements recorded at identical settings revealed that F-actin in WT cells was more or less evenly distributed around the entire phagocytic cup engulfing the yeast particles, whereas F-actin levels at the cup base of *forB*^*-*^ cells were moderately reduced by about 18% (Fig. 4*A* and *B*). Consistent with previous findings (39), *forG*^*-*^ cells exhibited a strong reduction in F-actin contents at the base of the cup of approximately 54%. Of note, the F-Actin content in the base of cups in *forB*^*-*^/*forG*^*-*^ cells was dramatically reduced by almost 70% on average, with many phagosomes completely lacking F-actin at the base (Fig. 4*A* and *B*). Moreover, we noticed that capturing of appropriate images for quantification of F-actin distribution in the double mutant proved exceptionally difficult, because these cells hardly formed any cups. Thus, we incubated WT and mutant cells at identical cell densities with the same amount of yeast particles in droplets on glass cover slips for 40 min prior to fixation, and then determined the fraction of cells containing yeast particles. WT, *forG*^*-*^ and *forB*^*-*^ cells internalized yeast particles at a similar rate, with about 50% of cells containing yeast particles (Fig. 4*C*). *ForB*^*-*^/*forG*^*-*^ cells, on the other hand, internalized yeast particles at a drastically reduced rate, with more than 77% of the double-mutant cells failing to internalize a single yeast particle. From these images we also determined the fraction of cells forming phagosomes. Approximately 12-15% of the WT cells and both single mutants had formed phagosomes at various stages of particle engulfment, whereas this value was drastically reduced to 2.4% in the *forB*^*-*^/*forG*^*-*^ double mutant (Fig. 4*D*). This was corroborated by confocal imaging and 3D reconstructions of phalloidin-stained WT and mutant cells in solution (Fig. 4*E*). Taken together these data strongly suggest that formin-generated actin filaments formed by ForB and ForG and accumulating at the cup base are decisive for the formation and integrity of phagocytic cups to allow for efficient engulfment of extracellular particles.

**Figure 4.**
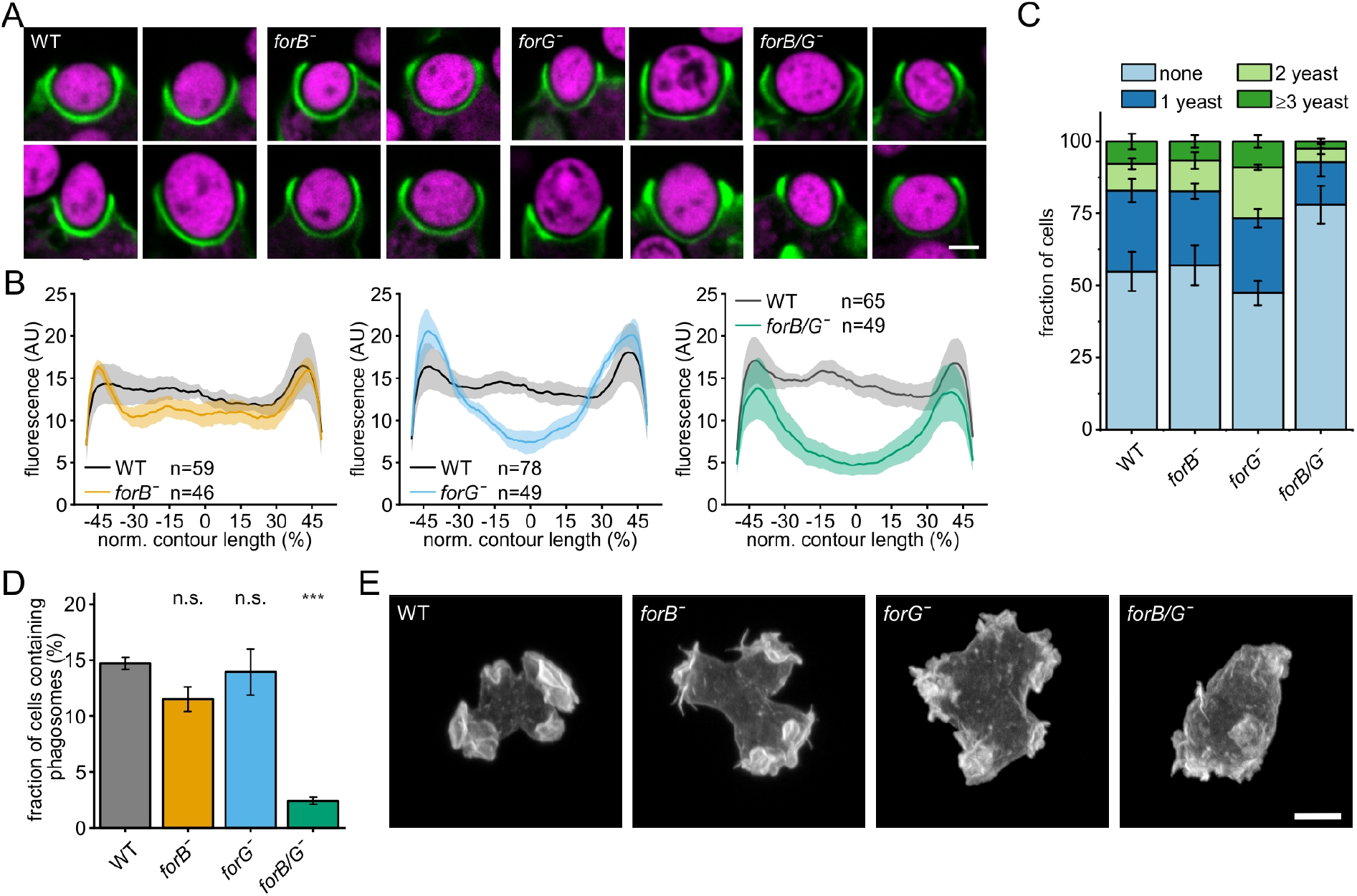
Combined loss of ForB and ForG severely perturbs phagosome formation. (*A*) Representative phagosomes of fixed WT and mutant cells during uptake of TRITC-labelled yeast particles (magenta). F-actin was stained with Atto488-phalloidin (green). Scale bar, 2 μm. (*B*) Quantification of the relative F-actin contents in the phagosomes of WT and mutant cells. Fluorescence intensities of the phalloidin-labeled F-actin structure along normalized contour lengths of phagosomes are shown (mean ± SD). The base of the phagosomes was set to 0%. AU, arbitrary units. Mean ± SD. n, number of analyzed cups. (*C*) Strongly impaired yeast phagocytosis of *forB*^*-*^/*forG*^*-*^ cells. Accumulation of TRITC-labeled yeast particles by phagocytosis after 40 min of incubation on glass coverslips. Confocal sections are shown. (*D*) Proportion of cells with phagosomes after 40 minute incubation with TRITC-labelled yeast particles followed by phalloidin staining. Note, *forB*^*-*^/*forG*^*-*^ double mutants had markedly fewer phagosomes compared to the WT or the other mutants. (*E*) Maximum intensity projections of WT and indicated mutant cells cultivated in growth medium and fixed and stained in solution for F-actin with phalloidin. Scale bar, 5 μm. Mean±SD, n=268-621 cells from 3-6 independent experiments. **p≤0.01, n.s. not significant.

### ForB promotes rocketing of internalized phagosomes

In time-lapse movies of cells phagocytosing yeast particles, we noticed that active YFP-ForBΔDAD gradually began to re-distribute from the cup base to its tip and then remained attached to the distal side of the phagosomes even after their closure and internalization. This was followed by the formation of a comet-like tail, pushing the yeast particle deeper into the cell (Fig. 5*A*, *SI Appendix*, Movie S5), thus strongly resembling the actin-driven rocketing of intracellular bacterial pathogens such as *Listeria* (46) or propelling beads in reconstituted motility assays (47). Co-expression of LifeAct-mRFP and YFP-ForB-N in WT cells confirmed co-localization of ForB in these actin comet tails (Fig. 5*B*, *SI Appendix*, Movie S6), and additionally excluded the possibility that localization was caused by excessive formin-driven actin assembly. Conversely, overexpression of active ForBΔDAD in WT cells led to a substantial increase in travelled distance and maximal speed of rocketing endosomes, whereas overexpression of active ForG did not cause similar effects (Fig. 5*C*-*E*). Finally, we analyzed rocketing parameters in the *forB*^-^/*forG*^*-*^double mutant. The combined loss of both formins led to a decrease in distance and maximal speed (reduced by 43% and 49%), but notably, these defects could be largely rescued by re-expression of active ForB, but interestingly not by expression of active ForG (Fig. 5*D* and *E*). Taken together, these data show that in spite of the overlapping functions of ForG and ForB in macroendocytosis, only ForB contributes to and promotes the rocketing of internalized phagosomes.

**Figure 5.**
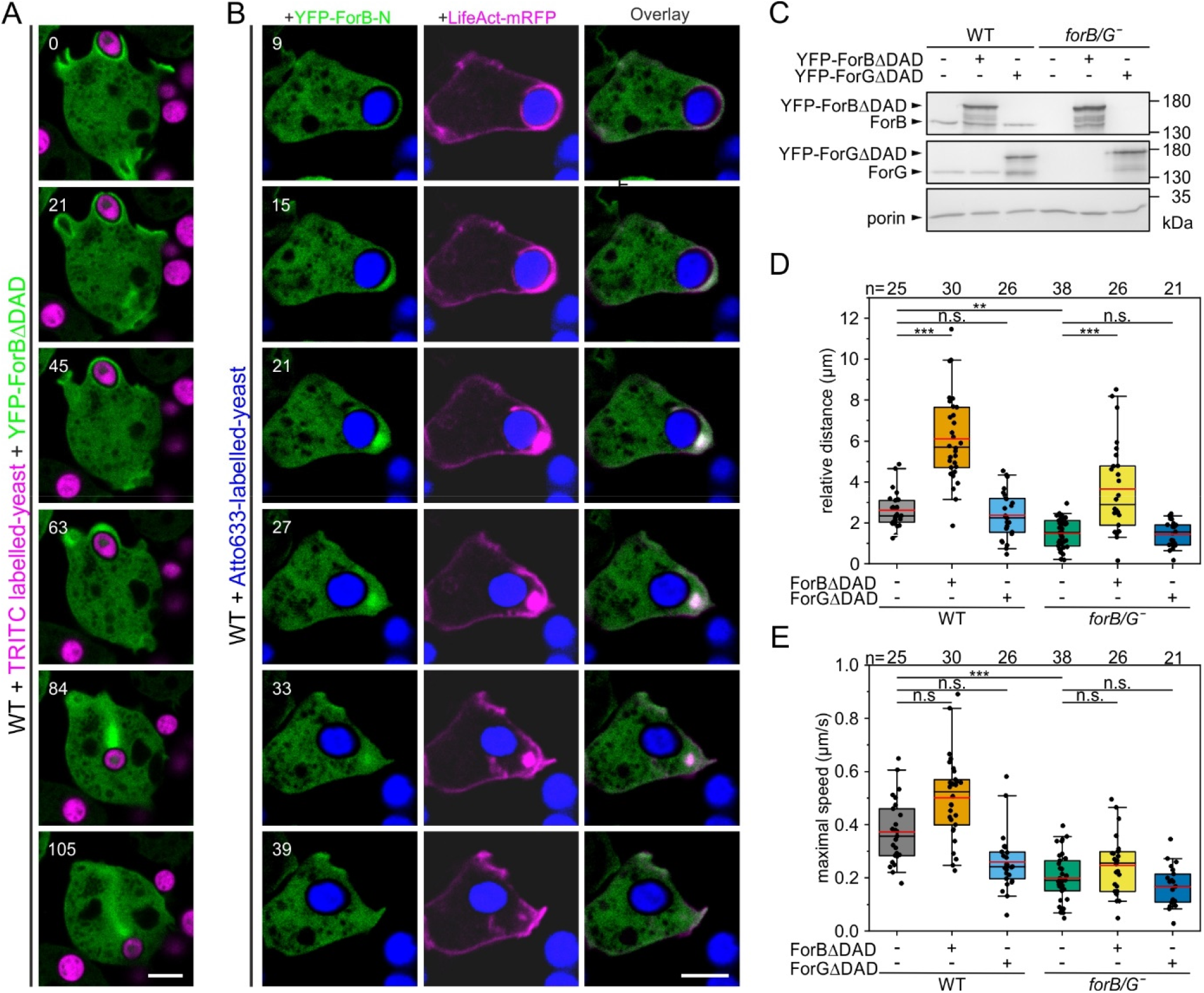
ForB promotes the rocketing of internalized phagosomes. (*A*) Still images from a confocal, time-lapse movie of WT cells expressing YFP-ForB-ΔDAD (green) during and after internalization of TRITC-labelled yeast particles (magenta). During particle internalization constitutively active ForB relocalized from the cup base to the distal side of the phagosomes promptly after cup closure, and formed a comet-like tail apparently pushing the particle deeper into the cell. (*B*) Confocal, time-lapse imaging of WT cells co-expressing YFP-ForB-N (green) and LifeAct-mRFP (magenta) during internalization of an Atto633-labelled yeast particle (blue). Note co-localization of ForB and F-actin (white areas in overlay) to comet tails of rocketing phagosomes. Scale bar, 5 μm. (*C*) Western blots showing overexpression or reconstitution of indicated formins in WT or *forB*^*-*^/*forG*^*-*^ cells used for analyses of phagosome rocketing. (*D, E*) Quantification of travelled distance and speed of rocketing phagosomes. Mean±SD, n=21-38 movies. ** p≤0.01, *** p≤0.001, n.s. not significant.

## Discussion

In the present study, we assessed the specific contribution of ForB to macroendocytosis in *D. discoideum* cells. Despite the clear localization of active ForB in macroendocytic cups and a moderately reduced F-actin content in the cup base upon its loss, unexpectedly, ForB-deficient single mutant cells were neither noticeably impaired in particle nor fluid uptake. However, this is consistent with its rather weak *in vitro* actin assembly activity and is moreover apparently due to the presence of the more potent formin ForG, which has been previously shown to be required for efficient macroendocytosis in *D. discoideum* cells (39). Thus, in accordance with their overlapping functions, both formins have to be eliminated simultaneously to almost completely suppress phagocytosis. Functional redundancy is a characteristic feature of many biological systems and is instrumental in safeguarding fundamental cellular processes. In line with this notion, functional networks allowing mutual complementation of certain actin-binding proteins have already been observed for the actin crosslinking proteins α-actinin and filamin (48) cortexillin I and II (49) as well as for cortical formins ForA, ForE and ForH in *Dictyostelium* or mDia1 and mDia3 in B16-F1 mouse melanoma cells (31).

Although ForB and ForG act synergistically in actin filament assembly in macroendocytic cups, it is likely that they also exhibit specific roles. Notably, the actin system of macrophages and *Dictyostelium* cells has the capacity to self-organize into waves that propagate on the planar, substrate-attached membrane of a cell (50). In a recent analysis of these actin waves, which resemble 2D-projections of frustrated phagocytic cups (3, 51, 52), active ForB showed a prominent localization to the transition area of the actin wave that is assumed to correspond to the tip of phagocytic cups, while it only weakly associated with the inner territory, which corresponds to the base of phagocytic cups and where active ForG was found (42). Thus, in accordance with the proposed homology between waves and phagocytic cups, ForB would have been expected to be localized more at the protrusive edge of phagocytic cups rather than the base. Interestingly, observed fluctuations of active ForB in the wave were not noticeably matched by an actin response, which is consistent with the weak actin assembly function for ForB found in this study. Instead these fluctuations appeared to be more related to deciding whether the wave should continue to propagate (42). This could also apply to phagocytic cup formation, and is supported by the finding that cells lacking ForB and ForG not only exhibit aberrant cup morphology but also appear to be impaired in the initiation of cup formation.

DRFs necessitate interaction with the active forms of small Rho-family GTPases with their GBD to release autoinhibition and mediate activation and subcellular targeting (53). While ForG is regulated by active Ras (39), ForB, as shown here, is clearly regulated by Rac. Our Y2H-analysis revealed that at intermediate stringency the ForB-GBD can interact with the active variants of the five Rac GTPases Rac1a-c, RacB and RacF1, all belonging to the subset of the classical Rac proteins exhibiting highest degree of sequence conservation with human Rac1 (44, 54). Notably, Rac1a-c were all shown to regulate macroendocytosis (55). RacF1 was also shown to associate with sites of macroendocytosis, albeit its individual loss did not impair uptake (56). Interestingly, the GBD of ForB exhibited strongest interaction with active RacB which has been previously identified a regulator of various cytoskeletal function including macroendocytosis, chemotaxis and morphogenesis (57, 58). Interestingly, active RacB was additionally found to interact with the IQGAP-related protein GAPA (59) and p21-activated kinase (Pak)-family members (60, 61). A fluorescent reporter for active Rac based on the CRIB domain of PakB (Pak-GBD) was also found to faithfully accumulate in macroendocytic cups including the tip (17), suggesting that besides activating ForB in the non-protrusive cup base, RacB may also contribute to actin assembly driven by the Arp2/3 complex downstream of SCAR/WAVE regulatory complex activation at the protrusive front of the cups. Thus, RacB is likely to be simultaneously involved in the activation of different downstream actin assembly factors in these intertwined signaling cascades, which would also explain why loss of RacB does not fully phenocopy the loss of ForB. Consistent with this notion, it is also conclusive to converge Ras and Rac signaling pathways leading to the activation of the formins ForB and ForG to optimally coordinate actin assembly at the non-protrusive base of the cup with actin assembly at the tip. This raises the interesting question of how this coordination could be spatially achieved. Recently, the multidomain protein RGBARG, containing lipid-binding, RhoGEF, and RasGAP domains, was shown to be critical for the internalization of geometrically distinct bacteria (62). RGBARG localizes prominently to the tips of macropinocytic and phagocytic cups, and was proposed to restrict Ras activity mainly to the base of the cup and amplify Rac activity at the tip. Yet, another RasGAP referred to IqgC accumulates in the entire cup and was shown to suppress Ras signaling during macroendocytosis (63), adding an additional layer of complexity. Nevertheless, active Ras and Rac are both found in the base of the cups as evidenced by localization of respective probes (17). One potential candidate specifically activating RacB in the base is RacGEF1, not only because it has been shown to reside in the cell cortex and activate RacB, but also due to the fact that loss of either protein is accompanied with a similar defect in development (58).

Yet, the question arises as to why the cells specifically utilize formins for the generation of the non-protrusive cup base while they use the Arp2/3 complex to nucleate the dendritic filament network of the protrusive tip. Interesting clues have emerged from the analysis of the actin-rich cell cortex which constitutes a thin viscoelastic sheet of bundled or cross-linked actin filaments, nonmuscle myosin II, and associated proteins beneath the plasma membrane (64). Even though both Arp2/3 complex and the formin mDia1 were shown to generate the bulk of cortical actin filaments in Hela cells (65), atomic force microscopy measurements revealed that the short and more dynamic Arp2/3-nucleated filaments do not play an important role in cortical mechanics despite their much higher abundance (66). In contrast, the considerably longer, formin-nucleated filaments are effective in distributing mechanical stress over several micrometers, thus ensuring integrity to the cell cortex (66). Based on these finding we hypothesize that the formin-generated cup base is critical in providing the mechanical support of the cup structure to balance protrusive forces at the tip and enable efficient pseudopod extension. Thus, due to the absence of these long filaments, the *forB*^-^/*forG*^*-*^double mutant is apparently unable to generate the required mechanic support of the structure and therefore mostly fails to form cups. This seems to be of particular importance for the internalization of large particles, as confirmed here by the massively impaired uptake of yeast cells in the double mutant.

As yet, phagosome rocketing in *Dictyostelium* cells and macrophages was thought to be driven by asymmetric assembly of a branched filament network initiated by the Arp2/3 complex as evidenced by its specific accumulation at the comet tail (67, 68). Efficient elongation of the filaments was further found to require the actin polymerase VASP and profilin (28, 68). Our live cell imaging and analysis of mutants internalizing yeast particles extends this repertoire of actin assembly factors with a formin promoting the nucleation and elongation of linear filaments as demonstrated by the condensation of active ForB at the distal side of the phagosome. Comparable to VASP (28), ForB is not essential for rocketing, but affects travelled distance and speed of propelling phagosomes. Intriguingly, even though ForB and ForG synergize in the formation of phagocytic cups and phagocytosis, it was surprising to find that the function in rocketing is conferred only by ForB. The molecular cause is currently unclear, but a domain swapping approach combined with quantitative analyses of phagosome rocketing in reconstituted *forB*^-^/*forG*^*-*^double mutant cells expressing formin chimeras might prove helpful in identifying the relevant region in ForB in future work.

## Materials and Methods

### Plasmids

The full-length coding sequence of Formin B (aa 1-1126, *Dictybase* accession number: DDB_G0282297) was amplified from a cDNA library using specific primers. Subsequently, the single internal *BglII* site was inactivated by introducing a silent mutant by site-directed PCR mutagenesis using standard procedures. The cDNA was then inserted into *BglII* and *SpeI* sites of the expression plasmid pDM334 (69). After sequence verification, this plasmid served as a template for all subsequent ForB constructs. The sequences encoding constitutively-active ForBΔDAD (aa 1-1081) and the N-terminal construct ForB-N (aa 1-406) were amplified by PCR and inserted into the *BglII* and *SpeI* sites of expression plasmid pDM304-YFP (39). For co-expression with mRFP-tagged LifeAct (70) the entire expression cassette was excised from pDM344-LifeAct-mRFP (39) with *NgoMIV* and inserted into the same site of pDM304-YFP harboring either ForBΔDAD or ForB-N expression cassettes, respectively. For co-expression of mRFP-LifeAct together with YFP-ForGΔDAD the bicistronic vector was generated accordingly, using previously described pDM304-YFP-ForGΔDAD (39, 71). Bicistronic expression plasmids for bifluorescence complementation assays were created as described (39). In short, the ForBΔDAD sequence was inserted into the *BglII* and *SpeI* sites of pDM304 (72), containing the C-terminal β-sheet of mVenus (aa 211-238) to yield pDM304-VC-ForBΔDAD, in which the mVenus fragment is fused to the N-terminus of ForBΔDAD. RacB sequences encoding either WT, constitutively-active RacB Q61L or dominant-negative RacB T17N were inserted into the *BglII/SpeI* sites of pDM344 containing the remaining part of mVenus (aa 1-210). To ensure comparable expression levels of RacB and ForB constructs, the mVenus-Rac expression cassettes of the pDM344 were excised with *NgoMIV* and inserted into the same site of pDM304-VC-ForBΔDAD. The *forB* targeting vector was constructed by amplification of a 0.5 kb 5′ *PstI/BamHI* fragment and a 0.5 kb 3′ *SalI/HindIII* fragment of the *forB* gene from genomic DNA following standard procedures. The fragments were successively inserted into the corresponding sites of pLPBLP (73). The *racB* targeting vector was constructed accordingly. The pDM vectors were obtained from Dicty Stock Center (74).

For expression of the ForB-FH1-FH2-C construct harboring two poly-proline stretches (ForB-2pP, aa 571-1126) in *E. coli*, the codon-optimized sequence was synthesized (BioCat) and inserted into the *BamHI* and *SalI* sites of pGEX-6P-1 (GE Healthcare). For expression of ForB-N (aa 1-406) corresponding sequences were amplified by PCR and inserted into the *BamHI* and *SalI* sites of pGEX-6P-1. The generation of ForG-3P construct has been described previously (39). All constructs were verified by DNA sequencing. Sequences of all oligonucleotides are provided in *SI Appendix* Table S1.

### Protein purification

GST-tagged proteins were expressed in *E. coli* host Rosetta 2 (Novagen) by induction with 0.75 mM isopropyl-β-D-thiogalactoside at 24 °C for 12 h. The proteins were then purified from bacterial extracts by affinity chromatography using glutathione-conjugated agarose (Macherey-Nagel), followed by cleavage of the GST-tag with PreScission protease. The GST-tag was removed by a second affinity chromatography step with glutathione-agarose and the proteins in the flow-through were further purified by size exclusion chromatography using an Äkta Purifier System equipped with a HiLoad 26/600 Superdex 200 column (GE Healthcare). Protein-containing fractions were pooled and dialyzed against storage buffer (30 mM HEPES, pH 7.4, 150 mM KCl, 1 mM DTT, 60% (v/v) glycerol) for storage of ForB-2pP at -20°C or against immunization buffer (150 mM NaCl, 25 mM Tris-HCl, pH 8.0) for generation of ForB-specific polyclonal antibodies. The purification of ForG-3pP has been described previously (39). Recombinant, tag-free *Dictyostelium* PFN I was purified by poly-L-proline affinity chromatography as described (27). Ca^2+^-ATP actin was purified from rabbit skeletal muscle as described (75), stored in G-buffer (5 mM Tris-HCl, pH 8.0, and 0.2 mM CaCl_2_, 0.5 mM DTT, 0.2 mM ATP), and labelled on Cys374 with Atto488-maleimide (ATTO-TEC) for TIRF-M imaging or with N-(1-Pyrenyl)maleimide (Invivogen) for pyrene-actin polymerization assays, respectively.

### Pyrene-actin polymerization assays

Bulk actin polymerization assays were performed in black, 96-well microtiter plates (BRAND GmbH) containing 40 μl of the respective protein of interest appropriately diluted in 1x KMEI (50 mM KCl, 1 mM MgCl_2_, 1 mM EGTA, 10 mM imidazole, pH 7.0) and 130 μl of 1.23x KMEI with 0.0615% antifoam solution (Extran AP33, Merck). Actin polymerization was initiated by injection of 30 μl of a 13.33 μM solution of 10% pyrene-labelled G-actin in G-buffer (5 mM Tris-HCl, pH 8.0, 0.2 mM ATP, 0.1 mM CaCl_2_, 0.5 mM DTT) into the protein mix using the automated dispenser of a Synergy 4 microplate reader (BioTek) to a final volume of 200 μl and a final concentration of 2 μM actin in 1x KMEI with 0.05% antifoam. After mixing for 2 s, pyrene fluorescence was monitored using the 364 nm excitation and 407 nm emission wavelengths at 20 s intervals for 30 min. Data were normalized and analyzed with Excel (Microsoft).

### TIRF-M

For total internal reflection fluorescence microscopy, TIRF-flow chambers were assembled from mPEG-Silane (MW 2000, Lysan Bio)-coated 24×24 mm glass coverslips (Menzel-Gläser) fixed to microscope slides by double-sided adhesive tape. The chambers were then pre-incubated with Pluronic solution, containing 1% Pluronic F-127 and 1% bovine serum albumin (BSA) in double-distilled H_2_O for 5 min and subsequently washed with 1x BSA/KMEI (1% BSA in 1x KMEI) buffer. The prepared reaction mix contained TIRF buffer (20 mM imidazole, pH 7.4, 50 mM KCl, 1 mM MgCl_2_, 1 mM EGTA, 0.2 mM ATP, 15 mM glucose, 20 mM β-mercaptoethanol, 0.25% (w/v) methylcellulose, 20 μg·ml^-1^ catalase and 100 μg·ml^-1^ glucose oxidase) and appropriate dilutions of the proteins of interest in 1x KMEI. The assay was initiated by addition of G-actin (0.6-1.0 μM final actin concentration containing 10% Atto488-labelled-actin) to the reaction mix, followed by thoroughly mixing and flushing the mixture into the prepared flow chambers. Images from a Nikon Eclipse TI-E inverted microscope equipped with a TIRF Apo N 60x objective were captured at 4 s intervals with an exposure time of 70 ms using an Ixon3 897 EMCCD camera (Andor) for at least 10 min. The elongation rates of filaments were measured by manual tracking of growing filament barbed ends using Fiji/ImageJ. The nucleation efficacies were determined by counting and averaging the number of formed actin filaments 10 minutes after initiation of the reaction in 3 independent movies.

### Antibodies and immunoblots

Polyclonal antibodies against ForB were raised by immunization of a female New Zealand White rabbit with recombinant ForB-N as antigen following standard procedures. ForG antibodies have been described (39). Immunoblotting was performed according to standard protocol, using undiluted hybridoma supernatants of contact site A-specific mAb 33-294-17 (76), porin-specific mAb 70-100-1 (77) or polyclonal ForB or ForG antibodies (1:1000). Primary antibodies in immunoblots were detected with secondary alkaline phosphatase-coupled anti-mouse or anti-rabbit IgG (1:5000, Dianova) and the blots visualized with 5-Bromo-4-chloro-3-indolyl phosphate (BCIP, Carl Roth). Uncropped scans of immunoblots are show in *SI Appendix* Fig. S3.

### Cell culture, transfection and gene disruption

*Dictyostelium discoideum* AX2-214 wild type (WT) cells and derived cell lines were cultivated at 21 °C in HL5-C medium containing glucose (Formedium) on polystyrene-coated petri dishes or in shaken suspension at 150 rpm. Transfections were performed with the Xcell gene pulser (BIO-RAD) using the preset protocol 4-2-6 for *Dictyostelium* cells. pLPBLP knockout vectors (73) were linearized via *BamHI/SalI* digestion before transfection. Transfected cells were selected 24 h after transfection with either 10 μg·mL^-1^ Blasticidin S or 10 μg·mL^-1^ G418 (both InvivoGen). Gene disruptions were confirmed either by immunoblotting (*forB*^*-*^ and *forG*^*-*^) or PCR (*racB*^*-*^). *ForG*^*-*^ (strain JFL110) cells have been described previously (39). Removal of the floxed Blasticidin S resistance cassette for subsequent knockouts was performed by transient expression of Cre recombinase according to standard procedures (73, 78). Quantification of growth on bacterial lawns or in shaken suspension was performed as described (28).

### Quantification of macroendocytosis

Quantitative assessment of macropinocytosis and phagocytosis as well as preparation of fluorescently labelled yeast particles was performed as recently described (28). The clearance of bacteria from shaken suspension by *Dictyostelium* cells was quantified as described (39).

### Fluorescence microscopy and imaging

Growth-phase cells expressing fluorescent fusion proteins or BiFC constructs were seeded onto 3.5-cm-diameter glass-bottom dishes (Ibidi), allowed to adhere for at least 1 h, washed with LoFlo-medium (Formedium) and incubated in LoFlo-Medium for at least 6 h. Confocal images were obtained with a Zeiss LSM980 confocal microscope equipped with an alpha PlnApo 63x/NA 1.46 oil immersion objective using the laser lines 488 nm, 514 nm, 561 nm and 639 nm. TRITC-or Atto633-labelled yeast particles were added 10-15 minutes before imaging. Data were processed using ImageJ/Fiji and CorelDraw software.

### Quantification of F-actin distribution in phagocytic cups

WT or mutant cells were seeded onto ethanol-cleaned glass cover-slips for 1 h prior to addition of TRITC-labelled yeast particles in growth medium. After 40 minutes, the cells were fixed with a solution containing 2% paraformaldehyde, 10 mM PIPES, 0.18% picric acid, pH 6.6, extensively washed with PBS/glycine (1x PBS, pH 7.4, 100 mM glycine) and permeabilized with 70% ethanol for 3 min. After extensive washing with PBS/glycine, the fixed cells were blocked with PBG containing 1x PBS, pH 7.4, supplemented with 0.5% BSA, 0.045% cold fish gelatin (Sigma), prior to labeling of F-actin with Atto488-conjugated phalloidin (AttoTec) for 4 h (1:250 dilution) at room temperature. Finally, the cells were washed three times with PBG and embedded in gelvatol mounting medium. Confocal sections of cells with partially engulfed yeast particles (50-80% engulfment) were recorded at identical settings at the Zeiss LSM980 confocal microscope equipped with an alpha PlnApo 63x/NA 1.46 oil objective. A four-pixel-wide segmented line was used in Fiji/ImageJ to obtain fluorescence intensity profiles of the F-actin signal along the contour of the cup. To account for differences in cup size and engulfment stage, the intensity profiles were resampled to 100 data points representing 100% of the contour length with the *resample* function of MATLAB (MathWorks, Signal Processing Toolbox) and a custom build macro. Data were processed with Excel and Origin software. Quantification of yeast particles per cell as well as the percentage of cells containing phagosomes were obtained from the same samples. After acquisition of confocal sections, the Z-stacks were analyzed for presence of cells engulfing yeast particles using Fiji/ImageJ software. The cell counter plug-in was used to determine the total number of cells and the number of cells with phagosomes. The latter could be easily identified by the presence of the F-actin-rich cups containing the yeast particles.

### Quantification of phagosome rocketing

Confocal, time-lapse movies of WT and mutant cells expressing LifeAct-mRFP and indicated YFP-tagged formin constructs that just finished internalization of an Atto633-labelled yeast particle before onset of rocketing were used for analysis. Travelled distance and speed of the rocketing phagosomes were determined by measuring the distance between the respective site of phagocytosis and the phagosome containing yeast particle in consecutive frames of the movies using Fiji/ImageJ. The distance and maximal speed were calculated with Excel (Microsoft).

### Y2H Assay

Yeast-two-hybrid analysis was performed with the MATCHMAKER GAL4 Two-Hybrid System 3 (Clontech) essentially as described (39). In short, ForB-N (aa 1-406) was inserted into the pGADT7 vector as prey using the *EcoRI* and *BamHI* restriction sites and the previously reported collection of Ras- and Rho-family GTPase (39) were used as bait. The latter library was extended with constitutive-active RacB Q61L and dominant-negative RacB-T17N variants, which were obtained by site-directed PCR mutagenesis using the RacB WT sequence as template. Both constructs were inserted into the *EcoRI/BamHI* restriction sites of the bait vector pGBKT7. The AH109 yeast strain was co-transfected with bait and prey vector, and grown on synthetic double-dropout agar lacking leucine and tryptophan according to the manufacturer’s instructions. Several colonies were collected and grown in synthetic dropout liquid media overnight and then subjected to dropout agar plates. Protein interactions were scored by growth of cells on agar plates for 4 d at 25 °C on triple-dropout (TD) or quadruple-dropout (QD) agar plates. TD plates lacked leucine, tryptophan, and histidine, and were supplemented with 3 mM 3-amino-1,2,4-triazole to suppress the leaky HIS3-reporter gene according to the manufacturer’s instructions. QD plates lacked leucine, tryptophan, histidine, and adenine for highest stringency.

## Statistical Analysis

Statistical analyses were performed using Origin (Origin-Lab, 2021). Statistical significance of normally distributed data (Shapiro-Wilk test) was determined with the two-tailed, unpaired Student’s t-test and in case of multiple comparisons ANOVA (Tukey test) was used. In case of non-normal distribution, the Kruskal-Wallis-Test followed by Dunn’s post hoc test were used. Statistical significances are reported as * p < 0.5, ** p < 0.01, *** p < 0.001 and n.s. as non-significant.

## Acknowledgments

We thank Annette Breskott for technical assistance. This work was supported by a grant of the Deutsche Forschungsgemeinschaft (DFG) to JF (FA330/13-1).

## References

1. A. Aderem, D. M. Underhill, Mechanisms of phagocytosis in macrophages. Annu Rev Immunol 17, 593–623 (1999).

2. J. A. Swanson, Shaping cups into phagosomes and macropinosomes. Nat Rev Mol Cell Biol 9, 639–649 (2008).

3. M. Rabinovitch, Professional and non-professional phagocytes: an introduction. Trends Cell Biol 5, 85–87 (1995).

4. J. J. Lim, S. Grinstein, Z. Roth, Diversity and Versatility of Phagocytosis: Roles in Innate Immunity, Tissue Remodeling, and Homeostasis. Front Cell Infect Microbiol 7, 191 (2017).

5. S. Arandjelovic, K. S. Ravichandran, Phagocytosis of apoptotic cells in homeostasis. Nat Immunol 16, 907–917 (2015).

6. J. D. Dunn, et al., Eat Prey, Live: Dictyostelium discoideum As a Model for Cell-Autonomous Defenses. Front Immunol 8, 1906 (2017).

7. W. F. Loomis, Dictyostelium discoideum, a developmental system (Academic Press Inc., New York, 1975).

8. J. S. King, R. R. Kay, The origins and evolution of macropinocytosis. Philos Trans R Soc Lond B Biol Sci 374, 20180158 (2019).

9. D. B. Mills, The origin of phagocytosis in Earth history. Interface Focus 10, 20200019 (2020).

10. N. Yutin, M. Y. Wolf, Y. I. Wolf, E. v Koonin, The origins of phagocytosis and eukaryogenesis. Biol Direct 4, 9 (2009).

11. J. Martín-González, J.-F. Montero-Bullón, J. Lacal, Dictyostelium discoideum as a non-mammalian biomedical model. Microb Biotechnol 14, 111–125 (2021).

12. M. J. Carnell, R. H. Insall, Actin on disease--studying the pathobiology of cell motility using Dictyostelium discoideum. Semin Cell Dev Biol 22, 82–88 (2011).

13. G. Bloomfield, et al., Neurofibromin controls macropinocytosis and phagocytosis in Dictyostelium. Elife 4 (2015).

14. R. Sussman, M. Sussman, Cultivation of Dictyostelium discoideum in axenic medium. Biochem Biophys Res Commun 29, 53–55 (1967).

15. J. H. Vines, J. S. King, The endocytic pathways of Dictyostelium discoideum. Int J Dev Biol 63, 461–471 (2019).

16. V. Jaumouillé, C. M. Waterman, Physical Constraints and Forces Involved in Phagocytosis. Front Immunol 11, 1097 (2020).

17. D. M. Veltman, et al., A plasma membrane template for macropinocytic cups. Elife 5 (2016).

18. S. A. Freeman, S. Grinstein, Phagocytosis: receptors, signal integration, and the cytoskeleton. Immunol Rev 262, 193–215 (2014).

19. E. Uribe-Querol, C. Rosales, Phagocytosis: Our Current Understanding of a Universal Biological Process. Front Immunol 11, 1066 (2020).

20. S. Cornillon, et al., An adhesion molecule in free-living Dictyostelium amoebae with integrin β features. EMBO Rep 7, 617–621 (2006).

21. M. Pan, X. Xu, Y. Chen, T. Jin, Identification of a Chemoattractant G-Protein-Coupled Receptor for Folic Acid that Controls Both Chemotaxis and Phagocytosis. Dev Cell 36, 428–439 (2016).

22. M. B. Hallett, An Introduction to Phagocytosis. Adv Exp Med Biol 1246, 1–7 (2020).

23. S. Buracco, S. Claydon, R. Insall, Control of actin dynamics during cell motility. F1000Res 8 (2019).

24. D. J. Seastone, et al., The WASp-like protein scar regulates macropinocytosis, phagocytosis and endosomal membrane flow in Dictyostelium. J Cell Sci 114, 2673–2683 (2001).

25. M. A. Wear, A. Yamashita, K. Kim, Y. Maéda, J. A. Cooper, How capping protein binds the barbed end of the actin filament. Curr Biol 13, 1531–1537 (2003).

26. J. Faix, K. Rottner, Ena/VASP proteins in cell edge protrusion, migration and adhesion. J Cell Sci 135 (2022).

27. D. Breitsprecher, et al., Clustering of VASP actively drives processive, WH2 domain-mediated actin filament elongation. EMBO J 27, 2943–2954 (2008).

28. S. Körber, J. Faix, VASP boosts protrusive activity of macroendocytic cups and drives phagosome rocketing after internalization. Eur J Cell Biol 101, 151200 (2022).

29. N. Courtemanche, Mechanisms of formin-mediated actin assembly and dynamics. Biophys Rev 10, 1553–1569 (2018).

30. J. Block, et al., FMNL2 drives actin-based protrusion and migration downstream of Cdc42. Curr Biol 22, 1005–1012 (2012).

31. C. Litschko, et al., Functional integrity of the contractile actin cortex is safeguarded by multiple Diaphanous-related formins. Proc Natl Acad Sci U S A 116, 3594–3603 (2019).

32. T. D. Pollard, Regulation of actin filament assembly by Arp2/3 complex and formins. Annu Rev Biophys Biomol Struct 36, 451–477 (2007).

33. T. Otomo, C. Otomo, D. R. Tomchick, M. Machius, M. K. Rosen, Structural basis of Rho GTPase-mediated activation of the formin mDia1. Mol Cell 18, 273–281 (2005).

34. P. Massol, P. Montcourrier, J. C. Guillemot, P. Chavrier, Fc receptor-mediated phagocytosis requires CDC42 and Rac1. EMBO J 17, 6219–6229 (1998).

35. A. D. Hoppe, J. A. Swanson, Cdc42, Rac1, and Rac2 display distinct patterns of activation during phagocytosis. Mol Biol Cell 15, 3509–3519 (2004).

36. D. J. Hackam, O. D. Rotstein, A. Schreiber, W. j Zhang, S. Grinstein, Rho is required for the initiation of calcium signaling and phagocytosis by Fcγ receptors in macrophages. J Exp Med 186, 955–966 (1997).

37. A. L. Bishop, A. Hall, Rho GTPases and their effector proteins. Biochem J 348 Pt 2, 241–255 (2000).

38. D. M. Veltman, M. G. Lemieux, D. A. Knecht, R. H. Insall, PIP_3_-dependent macropinocytosis is incompatible with chemotaxis. J Cell Biol 204, 497–505 (2014).

39. A. Junemann, et al., A Diaphanous-related formin links Ras signaling directly to actin assembly in macropinocytosis and phagocytosis. Proc Natl Acad Sci U S A 113, E7464–E7473 (2016).

40. F. Rivero, et al., A comparative sequence analysis reveals a common GBD/FH3-FH1-FH2-DAD architecture in formins from Dictyostelium, fungi and metazoa. BMC Genomics 6, 28 (2005).

41. C. Kitayama, T. Q. P. Uyeda, ForC, a novel type of formin family protein lacking an FH1 domain, is involved in multicellular development in Dictyostelium discoideum. J Cell Sci 116, 711–723 (2003).

42. M. Ecke, et al., Formins specify membrane patterns generated by propagating actin waves. Mol Biol Cell 31, 373–385 (2020).

43. H. Beug, F. E. Katz, G. Gerisch, Dynamics of antigenic membrane sites relating to cell aggregation in Dictyostelium discoideum. J Cell Biol 56, 647–658 (1973).

44. G. Vlahou, F. Rivero, Rho GTPase signaling in Dictyostelium discoideum: insights from the genome. Eur J Cell Biol 85, 947–959 (2006).

45. K. Ohashi, K. Mizuno, A novel pair of split venus fragments to detect protein-protein interactions by in vitro and in vivo bimolecular fluorescence complementation assays. Methods Mol Biol 1174, 247–262 (2014).

46. J. A. Theriot, T. J. Mitchison, L. G. Tilney, D. A. Portnoy, The rate of actin-based motility of intracellular Listeria monocytogenes equals the rate of actin polymerization. Nature 357, 257–260 (1992).

47. T. P. Loisel, R. Boujemaa, D. Pantaloni, M. F. Carlier, Reconstitution of actin-based motility of Listeria and Shigella using pure proteins. Nature 401, 613–616 (1999).

48. W. Witke, M. Schleicher, A. A. Noegel, Redundancy in the microfilament system: abnormal development of Dictyostelium cells lacking two F-actin cross-linking proteins. Cell 68, 53–62 (1992).

49. J. Faix, et al., Cortexillins, major determinants of cell shape and size, are actin-bundling proteins with a parallel coiled-coil tail. Cell 86, 631–642 (1996).

50. T. Bretschneider, et al., The three-dimensional dynamics of actin waves, a model of cytoskeletal self-organization. Biophys J 96, 2888–2900 (2009).

51. W. O. Fenn, The Adhesiveness of Leucocytes to Solid Surfaces. J Gen Physiol 5, 143–167 (1922).

52. G. Gerisch, Self-organizing actin waves that simulate phagocytic cup structures. PMC Biophys 3, 7 (2010).

53. A. Seth, C. Otomo, M. K. Rosen, Autoinhibition regulates cellular localization and actin assembly activity of the diaphanous-related formins FRLα and mDia1. J Cell Biol 174, 701–713 (2006).

54. F. Rivero, H. Dislich, G. Glöckner, A. A. Noegel, The Dictyostelium discoideum family of Rho-related proteins. Nucleic Acids Res 29, 1068–1079 (2001).

55. M. Dumontier, P. Höcht, U. Mintert, J. Faix, Rac1 GTPases control filopodia formation, cell motility, endocytosis, cytokinesis and development in Dictyostelium. J Cell Sci 113 (Pt 1), 2253–2265 (2000).

56. F. Rivero, et al., RacF1, a novel member of the Rho protein family in Dictyostelium discoideum, associates transiently with cell contact areas, macropinosomes, and phagosomes. Mol Biol Cell 10, 1205–1219 (1999).

57. E. Lee, D. J. Seastone, E. Harris, J. A. Cardelli, D. A. Knecht, RacB regulates cytoskeletal function in Dictyostelium spp. Eukaryot Cell 2, 474–485 (2003).

58. K. C. Park, et al., Rac regulation of chemotaxis and morphogenesis in Dictyostelium. EMBO J 23, 4177–4189 (2004).

59. S. Mondal, et al., Regulation of the actin cytoskeleton by an interaction of IQGAP related protein GAPA with filamin and cortexillin I. PLoS One 5, e15440 (2010).

60. S. Lee, et al., Dictyostelium PAKc is required for proper chemotaxis. Mol Biol Cell 15, 5456–5469 (2004).

61. M. de la Roche, A. Mahasneh, S.-F. Lee, F. Rivero, G. P. Côté, Cellular distribution and functions of wild-type and constitutively activated Dictyostelium PakB. Mol Biol Cell 16, 238–247 (2005).

62. C. M. Buckley, et al., Coordinated Ras and Rac Activity Shapes Macropinocytic Cups and Enables Phagocytosis of Geometrically Diverse Bacteria. Curr Biol 30, 2912–2926.e5 (2020).

63. M. Marinović, et al., IQGAP-related protein IqgC suppresses Ras signaling during large-scale endocytosis. Proc Natl Acad Sci U S A 116, 1289–1298 (2019).

64. P. Chugh, E. K. Paluch, The actin cortex at a glance. J Cell Sci 131 (2018).

65. M. Bovellan, et al., Cellular control of cortical actin nucleation. Curr Biol 24, 1628–1635 (2014).

66. M. Fritzsche, C. Erlenkämper, E. Moeendarbary, G. Charras, K. Kruse, Actin kinetics shapes cortical network structure and mechanics. Sci Adv 2, e1501337 (2016).

67. M. Clarke, A. Müller-Taubenberger, K. I. Anderson, U. Engel, G. Gerisch, Mechanically induced actin-mediated rocketing of phagosomes. Mol Biol Cell 17, 4866–4875 (2006).

68. F. S. Southwick, W. Li, F. Zhang, W. L. Zeile, D. L. Purich, Actin-based endosome and phagosome rocketing in macrophages: activation by the secretagogue antagonists lanthanum and zinc. Cell Motil Cytoskeleton 54, 41–55 (2003).

69. D. M. Veltman, I. Keizer-Gunnink, P. J. M. van Haastert, An extrachromosomal, inducible expression system for Dictyostelium discoideum. Plasmid 61, 119–125 (2009).

70. J. Riedl, et al., Lifeact: a versatile marker to visualize F-actin. Nat Methods 5, 605–607 (2008).

71. K. Ohashi, T. Kiuchi, K. Shoji, K. Sampei, K. Mizuno, Visualization of cofilin-actin and Ras-Raf interactions by bimolecular fluorescence complementation assays using a new pair of split Venus fragments. Biotechniques 52, 45–50 (2012).

72. D. M. Veltman, G. Akar, L. Bosgraaf, P. J. M. van Haastert, A new set of small, extrachromosomal expression vectors for Dictyostelium discoideum. Plasmid 61, 110–118 (2009).

73. A. R. Kimmel, J. Faix, Generation of multiple knockout mutants using the Cre-loxP system. Methods Mol Biol 346, 187–199 (2006).

74. P. Fey, R. J. Dodson, S. Basu, R. L. Chisholm, One stop shop for everything Dictyostelium: dictyBase and the Dicty Stock Center in 2012. Methods Mol Biol 983, 59–92 (2013).

75. J. A. Spudich, S. Watt, The regulation of rabbit skeletal muscle contraction. I. Biochemical studies of the interaction of the tropomyosin-troponin complex with actin and the proteolytic fragments of myosin. J Biol Chem 246, 4866–4871 (1971).

76. G. Bertholdt, J. Stadler, S. Bozzaro, B. Fichtner, G. Gerisch, Carbohydrate and other epitopes of the contact site A glycoprotein of Dictyostelium discoideum as characterized by monoclonal antibodies. Cell Differ 16, 187–202 (1985).

77. H. Troll, et al., Purification, functional characterization, and cDNA sequencing of mitochondrial porin from Dictyostelium discoideum. J Biol Chem 267, 21072–21079 (1992).

78. J. Linkner, B. Nordholz, A. Junemann, M. Winterhoff, J. Faix, Highly effective removal of floxed Blasticidin S resistance cassettes from Dictyostelium discoideum mutants by extrachromosomal expression of Cre. Eur J Cell Biol 91, 156–160 (2012).

