## Supplementary Information for "Convergence of Ras- and Rac-regulated formin pathways is pivotal for phagosome formation and particle uptake in *Dictyostelium*"

**Supplementary Figures**
**Figure S1**

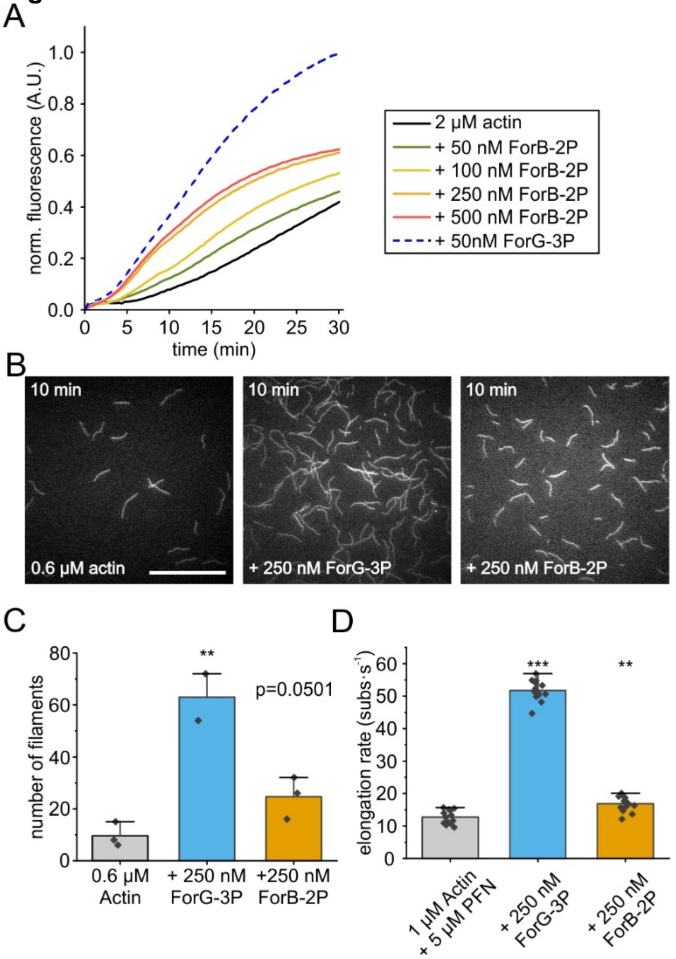


**Figure S1. ForB is a weaker actin assembly factor compared to ForG.** (*A*) ForB-2P promoted polymerization of pyrene-labelled G-actin (2 µM, 5% labeled) in a concentration-dependent manner, but was less efficient as compared to ForG-3P. A.U. arbitrary units. A.U. arbitrary units. (*B)* ForB is a weak nucleation factor compared to ForG-3P. Representative TIRF-M images after 10 minutes of actin-filament assembly are shown. Scale bar, 50 µm. (*C*) Quantification of the nucleation efficacies of ForB-2P *versus* ForG-3P in the absence of PFNI after 10 min. (*D*) Elongation rates of growing filaments as calculated from TIRF-M, time-lapse movies at indicated concentrations of ForG-3P and ForB-2P. At least 13 individual filaments from 3 independent movies were tracked. Mean±SD, ** p≤0.01, *** p≤0.001.

**Figure S2**

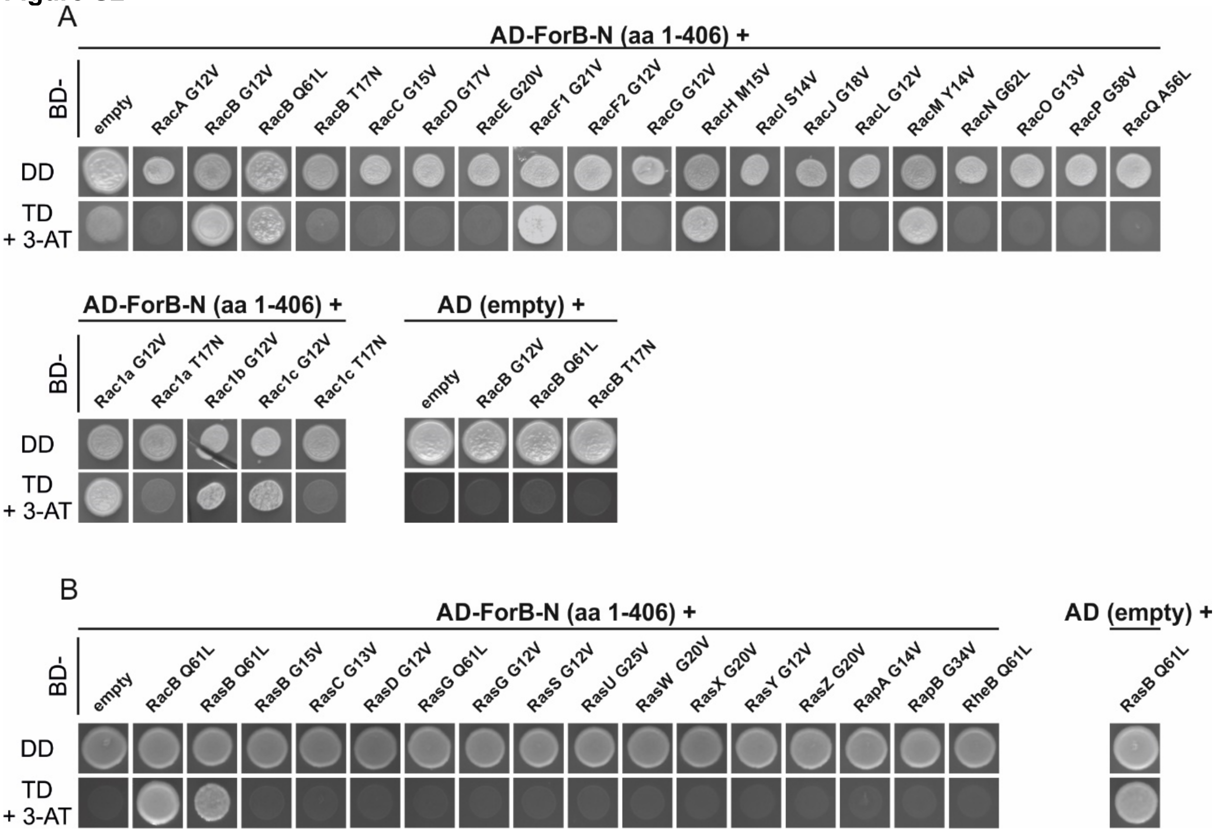


**Figure S2. ForB-N interacts with additional active *Dictyostelium* Rac-family GTPases under lower stringency.** (*A*) Y2H screen with constructs as shown in Fig. 4A at intermediate selection on triple dropout media (TD) supplemented with 3 mM 3-AT (+3-AT). Under these less stringent conditions, ForB interacted with additional Rac GTPases. (*B*) In contrast to ForG (Junemann et al., 2016), ForB did not interact with any of the tested active GTPases from the Ras family. The presence of prey and bait plasmids was monitored by growth on double dropout media (DD). RacB Q61L was used as positive control. Empty vectors were used as negative controls.

**Figure S3**

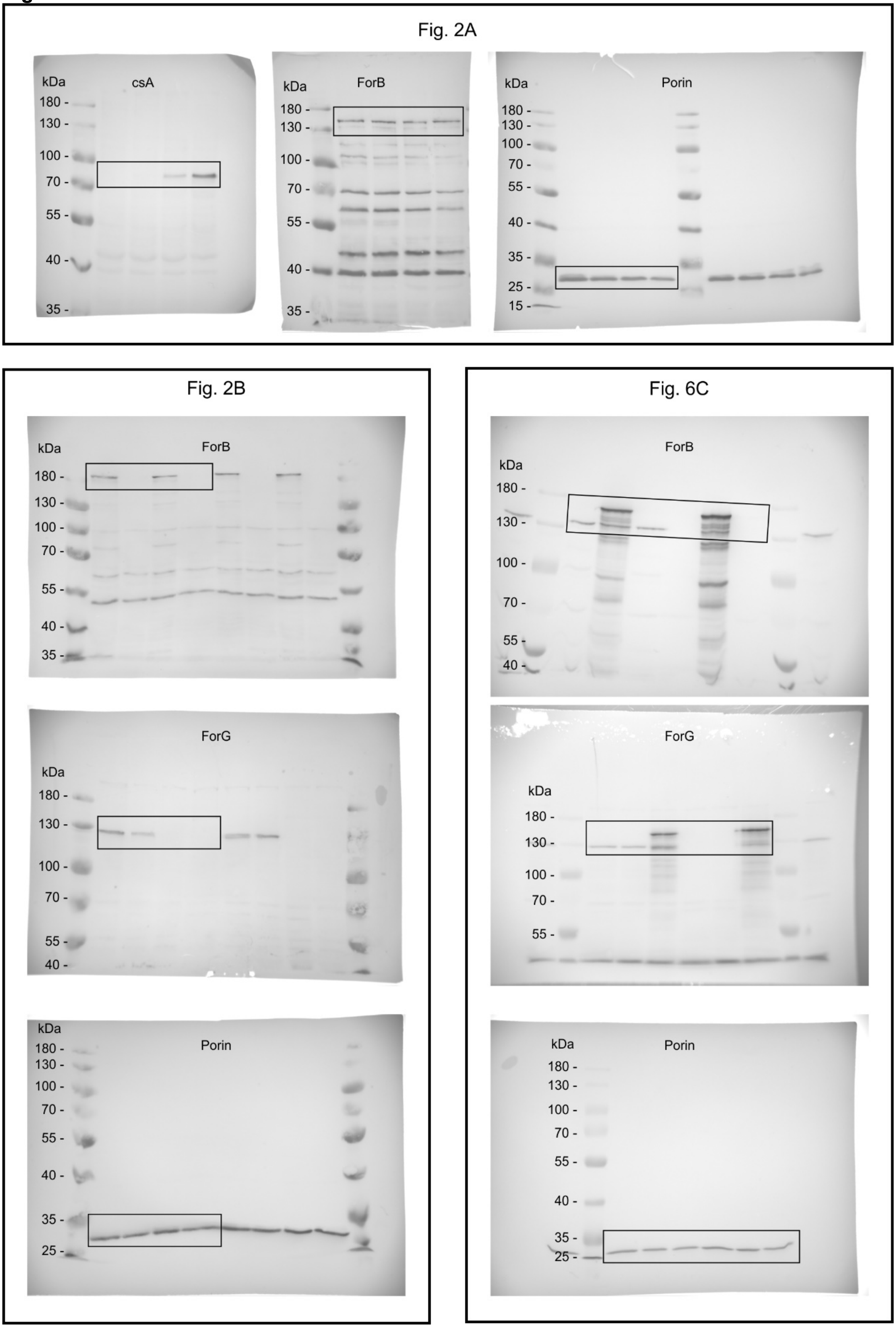


**Figure S3.** Uncropped images of immunoblots.

**Table**

**Table S1.** Sequences of oligonucleotides used in this work.

| Primer | Sequence | Orientation |
|---|---|---|
| *Dictyostelium* **constructs** | | |
| **GFP-ForB FL** | | |
| ForB FL Bgl fw | 5'-CGCAGATCTATGTTTTTTAAAGGTAAAAAAA-3' | forward |
| ForB FL Spe rev | 5'-GCGACTAGTTTATTTTTTATTTTTCAATGCAGC-3' | reverse |
| ForB FL Bam Mut rev | 5′-CGCGGATCCTGATATTGTTGTGGTGGAAG-3′ | reverse |
| **YFP-ForBΔDAD** | | |
| ForB-1081-ΔDAD-SpeD | 5'-CGCACTAGTATCCATAAAAGTACCATTTTGTGG-3' | reverse |
| **YFP-ForB-N** | | |
| ForB406SpeD-pDM | 5'-GCAACTAGTAGATTCTTTTGAAGGATCTTT-3' | reverse |
| ***forB*-KO** | | |
| ForB-BU-KO | 5'-GCGCGGATCCGCATGTTTTTTAAAGGTAAAAAAAAA-3' | forward |
| ForB-Pst-KO | 5'-GCGCTGCAGTAATGCTGCTGTCAATTGTAATACTG-3' | reverse |
| ForB-H3-KO | 5'-GCGAAGCTTTAATTCAATTCTTGGAGAATAAACATC-3' | forward |
| ForB-Sal-KO | 5'-CGCGTCGACTTCTAGATGCACGAATTGCTTCACCAC-3' | reverse |
| ***racB*-KO** | | |
| racB-5KO-BU | 5'-GCGGATCCGAGAAGAGAGGAATTAAATCT-3' | forward |
| racB-5KO-PD | 5'-GAGCTGCAGTACAGTTGGAACGTATTCTGT-3' | reverse |
| racB-3KO-HU | 5'-GAGAAGCTTGTCACACATCATTGTCCAAAC-3' | forward |
| racB-3KO-SD | 5'-CTCGTCGACACCATCATCACCATCGCAATC-3' | reverse |
| racB-KO-AD | 5'-CAATGATTCAACCAATGATTCAG-3' | reverse |
| Bsr-Koa | 5'-CAGTTACTCGTCCTATATACG-3' | forward |
| *E. coli* **constructs** | | |
| **ForB** | | |
| ForB-1-BU | 5'-TCAGGATCCATGTTTTTTAAGGTAAAAA-3' | forward |
| ForB-406-SD | 5'-CGCGTCGACTTAGAGATTCTTTTGAAGGAT-3' | reverse |
| ForB 2pP opt BglII | 5'-GATAGATCTAGCAGCGGCGGAGGC-3' | forward |
| ForB-SD | 5'-CTCGTCGACTTATTTTTTATTTTTCAATGCAGC-3' | reverse |
| **Y2H constructs** | | |
| ForB1EcoU-pGAD | 5'-CGCGAATTCATGTTTTTTAAAGGTAAAAAAAA-3' | forward |
| corForB406BD-pGAD | 5'-CTAGGATCCTTAAGATTCTTTTGAAGGATCTT-3' | reverse |
| RacB Q61L opt Eco fw | 5'-GTAGAATTCATGCAGTCCATCAAACTGGT-3' | forward |
| RacBQ61L opt dCAAX rev Bam | 5'-CGCGGATCCCTTGGAGTTTTTTTGTTGGTCGC-3' | reverse |

**Movie legends:**

**Movie S1**

ForB co-localizes with F-actin at sites of macropinocytosis. Confocal, time-lapse imaging of a WT cell expressing YFP-ForBΔDAD (green) and mRFP-LifeAct (magenta). Still from this movie are shown in Fig. 1D. Time is in min:s. Scale bar, 5 μm.

**Movie S2**

ForB localizes prominently to phagocytic cups. Confocal, time-lapse imaging of a WT cell expressing YFP-ForBΔDAD (green) during internalization of a TRITC-labelled yeast particle (magenta). Stills from this movie are shown in Fig. 1E. Time is in min:s. Scale bar, 5 μm.

**Movie S3**

ForB interacts with active RacB interact *in vivo* and localize to sites of phagocytosis. Confocal, time-lapse imaging of a WT cell expressing VC-ForBΔDAD and VN-RacB Q61L (green), during internalization of a TRITC-labelled yeast particle (magenta). Stills from this movie are shown in Fig. 4C. Time is in min:s. Scale bar, 5 μm.

**Movie S4**

ForB and active RacB interact *in vivo* and co-localize to macropinocytic cups. Confocal, time-lapse imaging of a WT cell expressing VC-ForBΔDAD and VN-RacB Q61L (green). Stills from this movie are shown in Fig. 4D. Time is in min:s. Scale bar, 5 μm.

**Movie S5**

Overexpression of active ForB promotes the rocketing of internalized phagosomes. Confocal, time-lapse imaging of a WT cell expressing YFP-ForBΔDAD (green) during and after internalization of a TRITC-labelled yeast particle (magenta). Stills from this movie are shown in Fig. 6A. Time is in min:s. Scale bar, 5 μm.

**Movie S6**

Confocal, time-lapse imaging of a WT cell expressing YFP-ForB-N (green) and mRFP-LifeAct (magenta) during internalization of an Atto633-labelled yeast particle (blue). Stills from this movie are shown in Fig. 6B. Time is in min:s. Scale bar, 5 μm.
